# Group-walk, a rigorous approach to group-wise false discovery rate analysis by target-decoy competition

**DOI:** 10.1101/2022.01.30.478144

**Authors:** Jack Freestone, Temana Short, William Stafford Noble, Uri Keich

**Affiliations:** School of Mathematics and Statistics F07 University of Sydney; Departments of Genome Sciences and of Computer Science and Engineering University of Washington

## Abstract

Target-decoy competition (TDC) is a commonly used method for false discovery rate (FDR) control in the analysis of tandem mass spectrometry data. This type of competition-based FDR control has recently gained significant popularity in other fields after Barber and Candès laid its theoretical foundation in a more general setting that included the feature selection problem. In both cases, the competition is based on a head-to-head comparison between an (observed) target score and a corresponding decoy (knockoff) score. However, the effectiveness of TDC depends on whether the data is homogeneous, which is often not the case: in many settings, the data consists of groups with different score profiles or different proportions of true nulls. In such cases, applying TDC while ignoring the group structure often yields imbalanced lists of discoveries, where some groups might include relatively many false discoveries and other groups include relatively very few. On the other hand, as we show, the alternative approach of applying TDC separately to each group does not rigorously control the FDR.

We developed *Group-walk*, a procedure that controls the FDR in the target-decoy / knockoff setting while taking into account a given group structure. Group-walk is derived from the recently developed AdaPT — a general framework for controlling the FDR with side-information. We show using simulated and real datasets that when the data naturally divides into groups with different characteristics Group-walk can deliver consistent power gains that in some cases are substantial. These groupings include the precursor charge state (4% more discovered peptides at 1% FDR threshold), the peptide length (3.6% increase) and the mass difference due to modifications (26% increase).

Group-walk is available at https://cran.r-project.org/web/packages/groupwalk/index.html

## 1 Introduction

Tandem mass spectrometry (MS/MS) currently provides the most efficient means of studying proteins in a high-throughput fashion. In a “shotgun proteomics” MS/MS experiment, proteins in a complex biological sample are extracted and digested into peptides, each with an associated charge. These charged peptides, called “precursors”, are measured by the mass spectrometer, and a subset of the precursors are then selected for further fragmentation into charged ions, which are detected and recorded by a second round of mass spectrometry [16, 26]. The resulting tandem fragmentation spectra, or spectra for short, are then subjected to computational analysis. A typical 30-minute MS/MS experiment will generate approximately 18,000 such spectra. Canonically, each observed spectrum is generated by a single peptide. Thus, the first goals of the downstream analysis are to identify which peptide generated each of the observed spectra (the spectrum-ID problem) and to determine which peptides and which proteins were present in the sample (the peptide/protein detection problems).

In each of those three problems the canonical approach to determining the list of discoveries is by controlling the false discovery rate (FDR) through some form of target-decoy competition (TDC). TDC has been widely practiced by the computational mass spectrometry community since it was first proposed by Elias and Gygi [9, 6, 17, 10, 14, 31].

To understand how TDC is applied in practice, consider for example the spectrum-ID problem. This analysis is typically initiated by using a search engine to scan each input spectrum against a target peptide database for its best matching peptide (in practice the spectrum is matched only against a subset of candidate peptides whose mass is within the measurement tolerance of the so-called precursor mass associated with the spectrum). Pioneered by SEQUEST [12], the search engine uses an elaborate score function to quantify the quality of the match between each of the database peptides and the observed spectrum, recording the optimal peptide-spectrum match (PSM) for the given spectrum along with its score *Z_i_* [25]. In practice, many expected fragment ions will fail to be observed for any given spectrum, and the spectrum is also likely to contain a variety of additional, unexplained peaks [26]. Hence, sometimes the reported PSM is correct — the peptide assigned to the spectrum was present in the mass spectrometer when the spectrum was generated — and sometimes the PSM is incorrect. Ideally, we would report only the correct PSMs, but obviously we do not know which PSMs are correct and which are incorrect; all we have is the score of the PSM, indicating its quality. TDC allows us to report a thresholded list of top-scoring PSMs while controlling the list’s FDR, as explained next.

First, the same search engine is used to assign each input spectrum a *decoy* PSM score, 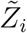, by searching for the spectrum’s best match in a decoy database of peptides obtained from the original database by randomly shuffling or reversing each peptide in the database. Each decoy score 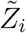 then directly competes with its corresponding target score *Z_i_* to determine the reported list of discoveries, i.e., we only report target PSMs that win their competition: 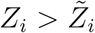. Additionally, the number of decoy wins 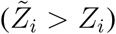 in the top *k* scoring PSMs is used to estimate the number of false discoveries in the target wins among the same top *k* PSMs. Thus, the ratio between the number of decoy wins and the number of target wins yields an estimate of the FDR among the target wins in the top *k* PSMs. To control the FDR at level *α*, the TDC procedure chooses the largest *k* for which the estimated FDR is still ≤ *α*, and it reports all target wins among those top *k* PSMs. It was recently shown that, assuming that *incorrect PSMs are independently equally likely to come from a target or a decoy match,* and provided we add 1 to the number of decoy wins before dividing by the number of target wins, this procedure rigorously controls the FDR [15].

As first noted by Efron in the context of the canonical p-value-based FDR analysis, the implicit assumption that FDR control relies on is that the data is essentially homogeneous. When, as often happens, the data is made of groups of hypotheses with distinct group-wise characteristics (e.g., different proportion of true nulls), performing a global FDR analysis can be problematic [8].

Efron’s observation is even more important in the context of TDC, where the issue of score calibration is known to play a significant role [17, 20]. For example, Baker, Medzihradszky, and Chalkley noted that spectra characteristics vary with the precursor’s charge state [1]. Specifically, they noted that by separately applying TDC to each charge state they obtain more discoveries at the 1% FDR threshold than they get when applying TDC to the combined dataset [1, Table V]. Arguably even more disturbing is the fact that, because their group-specific 1% FDR thresholds vary significantly [1, Table IV], inevitably when a single cutoff is used for the entire set, the discoveries from some groups will be overly conservative while for others they will be overly liberal.

Baker et al. tried to address the specific charge-state structure by redesigning their score function so it is better calibrated with respect to charge state. However, they also noted a similar problem of group-wise imbalance when considering phosphopeptides vs. unmodified ones: ignoring the distinction between the two groups meant that, in their case, the FDR in the phosphopeptides group was significantly underestimated [1, Table VI]. Their suggestion in this case coincided with Efron’s solution, which is to apply a separate FDR analysis to each group. Clearly, the latter is more attractive as it does not require us to redesign our score function to address any potential group structure. However, as Efron noted, applying a separate FDR analysis to each group can fail to control the FDR in the finite sample setting. Indeed, we show here that separately applying TDC to each group can significantly underestimate the true FDR.

Instead, we offer Group-walk a new procedure that shares the essence of separate TDC analyses but rigorously controls the FDR even for finite samples. Group-walk derives its theoretical guarantees from a recently published paper that developed a general framework for the problem of multiple hypothesis testing with generic side information; i.e., associated with each hypothesis is a p-value, as well as a predictor encoding contextual information about the hypothesis [22]. Of course, a group ID, as in our case, is a special case of such side information, but to the best of our knowledge we are the first to effectively adapt the general framework of Lei and Fithian to the context of group-wise TDC analysis.

We demonstrate the potential power of Group-walk using simulated as well as real datasets. The simulated data allows us to examine how Group-walk takes advantage of the different ways in which groups may vary. We complement the simulations with investigating multiple, naturally defined group structures in randomly selected real datasets. Specifically, we compare the number of peptides discovered from mass spectrometry data using TDC (ignoring the group structure) and using Group-walk, taking into account the charge state, peptide length and the mass difference due to post-translational modifications. In those analyses Group-walk overall delivers more discoveries across the entire range of examined FDR thresholds, varying, depending on the context, from a median increase of 3.6% all the way to a median increase of 26%.

Our motivation and our demonstrations are focused on the analysis of mass spectrometry data. However, Group-walk is in principle applicable in the broader context of competition-based approach to FDR, including its application to variable selection proposed by Barber and Candès [2]: if the data (variables) naturally partitions into groups with different characteristic the rigorous group-wise FDR analysis that Group-walk offers can be beneficial.

## 2 Background

The TDC procedure uses competition to control the FDR. The approach can be phrased in more general terms as follows. Let *H_i_* (*i* = 1,…, *m*) denote our *m* null hypotheses, e.g., in the spectrum-ID problem *H_i_* is “the ith PSM is incorrect” and in the peptide detection problem *H_i_* is “the ith peptide is present in the sample.” Associated with each *H_i_* are two competing scores: a target score *Z_i_* (the higher the score the less likely *H_i_* is) and a decoy score 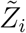. For example, in spectrum-ID *Z_i_* 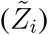 is the score of the optimal target (decoy) peptide match to the ith spectrum, whereas in peptide detection *Z_i_* corresponds to the maximal score of all PSMs that are optimally matched to the ith target peptide (and 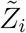 is similarly defined for the ith decoy peptide).

Adopting the notation of [11] we associate with each hypothesis a “winning” score *W_i_* and a target/decoy-win label *L_i_*. For the problems we consider here 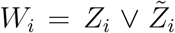 (where *x* V *y* = max{*x, y*}), but other functions can be considered as well [2]. As for *L_i_*, we define it as 1 if *H_i_* corresponds to a target win and as −1 if *H_i_* corresponds to a decoy win. Here we assume that ties 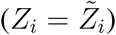 are broken randomly, but ignoring hypotheses with ties is also valid.

Let *N* = {*i*: *H_i_* is a true null} and note that while typically in the context of hypotheses testing *N* is a constant, albeit unknown set, here we allow *N* to be a random set. The fundamental assumption that TDC relies on is

### Assumption 1.

*Conditional on all the scores {W_i_}_i_ and all the false null labels {L_i_: i ∈ N}, the true nulls are independently equally likely to be a target or a decoy win, i.e., the random variables (RVs) {L_i_: i ∈ N} are conditionally independent uniform ±1 RVs.*

Clearly, if the assumption holds then {*L_i_*: *i* ∈ *N*} are still independent uniform ±1 RVs after ordering the hypotheses in increasing order (ties broken randomly) so that *W*_1_ ≤ *W*_2_ ≤ … ≤ *W_m_*. Hence, without loss of generality the scores are assumed to be in increasing order.

Let 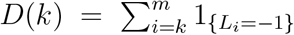 (the number of decoy wins with rank ≥ *k*), and let 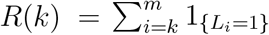 (the corresponding number of target wins). TDC defines its discovery list as {*i* ≥ *K*: *L_i_* = 1}, i.e., all the target wins with rank ≥ *K*, where *K = K*(*α*) = min{*k*: (*D*(*k*) + 1)/(*R*(*k*) ∨ 1) ≤ *α*} is TDC’s rejection threshold.

It was proved by He et al. that TDC controls the FDR in the finite sample setting [15]. That is, with 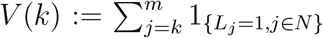 (the number of true null target wins with rank ≥ *k*), the FDP among the corresponding target wins is *Q_k_* = *V*(*k*)/(*R*(*k*) ∨1), and *E*(*Q_K_*) ≤ *α*. The expectation is taken with respect to the labels of the true null hypotheses conditional on the false null labels and on all the given scores.

At essentially the same time Barber and Candès established the same result through a sweeping generalization that deals with sequential hypothesis testing. In that setup the order of the hypotheses is predetermined, and a p-value is associated with each hypothesis so that given the order, as well as the p-values of the false nulls, the p-values of the true null hypotheses are indendent and identically distributed with a distribution that stochastically dominates the uniform distribution (so they are valid p-values). In our case we order the hypotheses by the scores *W_i_*, and the p-values are defined as a “1-bit p-value”: 1 for a decoy win and 1/2 for a target win. It is easy to see that Assumption 1 guarantees that the conditions of the sequential hypothesis testing are satisfied so it indeed generalizes TDC’s setup.

One reason TDC was adopted in the mass spectrometry context, rather than relying on standard methods for control of the FDR such as the procedures by Benjamini and Hochberg [3] or Storey [29], is that the latter require sufficiently informative p-values and, initially, no such p-values were computed in this context (as noted above, using the decoys we can always assign a 1-bit p-value to the hypotheses, but those are not informative enough to obtain effective results using the latter procedures). Moreover, the proteomics dataset typically consist of both “native” spectra (those for which their generating peptide is in the target database) and “foreign” spectra (those for which it is not). These two types of spectra create different types of false positives, implying that we typically cannot apply the standard FDR controlling procedures to the spectrum-ID problem even if we are able to compute p-values [18].

## 3 Methods

### 3.1 Group-walk

Group-walk is our novel procedure designed to analyze the same target-decoy data that TDC does while taking advantage of the data’s group structure. Specifically, in addition to the label *L_i_* and the score *W_i_*, each hypothesis has a group identifier *G_i_* ∈ {1, 2,…, *n_g_*} where *n_g_* ≥ 1 is the number of groups. Accordingly, we revise Assumption 1 to account for the group information:

#### Assumption 2.

*Conditional on all the scores {W_i_}_i_, the group IDs {G_i_}_i_ and all the false null labels {L_i_: i ∈ N}, the true nulls are independently equally likely to be a target or a decoy win.*

To understand how Group-walk works it is instructive to revisit TDC. Assuming the scores W_i_ are in increasing order, we define the “front” as the index *k* that we are currently considering as the rejection threshold. Starting with the front set to *k* = 1, TDC iteratively checks if the FDR estimated from all the hypotheses that the front is yet to pass over is below the threshold *α* (i.e., if (*D*(*k*) + 1)/(*R*(*k*) ∨1) ≤ *α*). If it is, then TDC stops and reports the location of the front (implying that all the target-winning hypotheses that the front is yet to pass over are rejected, or considered discoveries); otherwise, TDC advances the front by one step *k*: = *k* + 1.

TDC considers a single group, and therefore a 1-dimensional front, whereas Group-walk’s front is an *n_g_*-dimensional vector **k** = (*k_b_*…, *k_n_g__*), where each *k_g_* ∈ {1, 2,…, *m_g_*} and *m_g_* is the size of the *g*th group. Assuming the scores 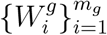 in each group are sorted in increasing order (with corresponding labels 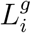), Group-walk starts with the trivial front **k** = (1,…, 1), i.e., the first index (lowest scoring hypothesis) in each of the groups. Just like TDC, it then iteratively checks if the FDR estimated from all the hypotheses that the front is yet to pass over is below the threshold *α*. For Group-walk this translates to checking whether 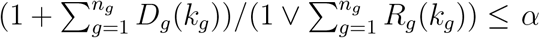, where 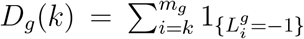 (the number of decoy wins in group *g* with rank ≥ *k*) and 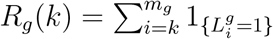 (the corresponding number of target wins).

If that estimated FDR is ≤ α, then Group-walk stops and returns its current front (implying all the target-winning hypotheses that the front is yet to pass over are rejected); otherwise, Group-walk advances the front by one step along a single coordinate. This is where Group-walk fundamentally differs from TDC: its front is *n_g_*-dimensional so it needs to choose which of the *g* groups should be advanced.

Group-walk employs an initial trivial strategy of iteratively advancing one group at a time until the front reaches a pre-determined index *K* in each group: **k** = (*K, K*,…, *K*) (here we used *K* = 40). Note that in some rare cases Group-walk might stop before the front reaches this point, and if the front needs to advance beyond the length of a group then that group is tossed away with no discoveries.

When Group-walk reaches the uniform front of **k** = (*K, K,…, K*) it switches its strategy to try to predict which of the groups is most likely to have a decoy at the current location of the front. That is, considering 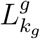 for *g* = 1,…, *n_g_*, which one is most likely = — 1? The motivation for this approach is that any hypothesis that the front passes over is “lost”: it can no longer be discovered. Of course, hypotheses for which 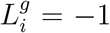 cannot be discoveries because they are decoy wins, so we might as well sacrifice those. Another way of looking at it is that where the chances of 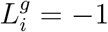 are higher it is less likely be a region of high quality target wins.

Group-walk attempts to solve the question of which 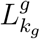 is most likely = – 1 by looking at the last K labels of each group and choosing the one that has the most decoy wins. The algorithmic description of Group-walk is provided in Algorithm 1 in the Supplement.

In Supplementary Section 6.4 we provide the details establishing that Group-walk controls the FDR in the finite-sample setting by showing it is a special case of (a slightly modified version of) the Adaptive p-value Thresholding (AdaPT) framework of Lei and Fithian [22]. AdaPT was developed to address the problem of controlling the FDR, where associated with each hypothesis is a p-value *p_i_* as well as a predictor *x_i_* encoding contextual information about the hypothesis. In our case the p-value is the 1-bit p-value and *x_i_* = (*W_i_, G_i_*).

### 3.2 Simulations

Groups of hypotheses can roughly differ along the following three attributes that might vary between the groups: the distribution of the null scores (“group null-effect”), the degree of separation between the true and false null scores (“group separation-effect”), and the proportion of false nulls (“group π-effect”). We use simulated data to look at how Group-walk exploits those effects by applying it to a group structure of two groups that differ with respect to some of those attributes while agreeing on the others. Additionally, we looked at how differences in group sizes can accentuate or moderate group effects.

We focused our simulations on the spectrum-ID problem using a previously published model [19, 18]. Briefly, each virtual “spectrum” *σ_i_* is associated with three randomly drawn scores: the score *X_i_* of the match between *σ_i_* and its generating peptide, the score *Y_i_* of the best match to *σ_i_* in the target database minus the generating peptide (if *σ_i_* is native), and the score 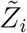 of the best match to *σ_i_* in the decoy database. The three scores are drawn independently of one another as well as of the corresponding scores of all other spectra. More specifically, *Y_i_* and 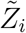 are sampled from a null distribution (which can be spectrum-specific), whereas *X_i_* is sampled from an alternative distribution for a native *σ_i_*, and *X_i_* is set to the lowest possible score for a foreign *σ_i_*. The target PSM score is *Z_i_* = *X_i_* ∨ *Y_i_*, and the decoy PSM score is 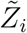. Finally, the PSM is incorrect when 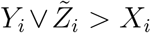. It is easy to see that conditional on the PSM being incorrect, 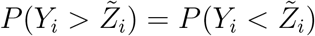 independently of everything else; hence, the spectrum-ID model is indeed captured by Assumption 1.

In Supplementary Section 6.1 we describe in detail how given the sizes *m*_1_, *m*_2_ ∈ {1e4, 2e4, 4e4} of the two groups of null hypotheses (“the PSM is incorrect, i.e., 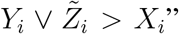), the separation parameters *b*_1_,*b*_2_ ∈ {5,50}, and the proportions of native spectra (which are related to but not identical to the proportion of false nulls, cf. Section 2) *π*_1_, *π*_2_ ∈ {0.3,0.6, 0.9} we draw a random dataset of two groups of scores with associated target/decoy win labels.

A score is “calibrated” if all true null scores are generated according to the same distribution. Hence, in order to examine a group null-effect we simulated uncalibrated scores. Specifically we simulated a group null-effect due to distinct precursor charge states where the first group simulates PSMs with a precursor charge state 3 and the second group simulates charge 2 (see Supplementary Section 6.1 for details).

To smooth out random effects due to the random nature of our draws we drew 40 random datasets (“runs”) for each of the 628 parameter combinations we considered. We then applied Group-walk (with *K* = 40), as well as TDC to each drawn dataset, where TDC was applied while ignoring the group structure, and the ratio of discovery numbers was averaged over the 40 runs.

### 3.3 Analysis of mass spectra

To test Group-walk on real data sets we randomly sampled one spectrum file from each of 14 randomly selected projects from the Proteomics Identifications Database (PRIDE) [28]: 7 high-resolution and 7 low-resolution spectra files. We then used the Crux toolkit [27] to (a) create a randomly shuffled decoy database from each project’s target peptide database, and (b) generate the PSMs by searching each spectrum against the concatenated target-decoy peptide database (for details see Supplementary Section 6.2).

When analyzing real data the spectrum-ID analysis can be problematic because the same peptide can generate multiple spectra, which can potentially break the true-nulls independence assumption and violate FDR control [15]. Thus, we chose to focus on the peptide-level analysis of the real data. In our peptide-level analysis the score 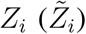 we assign to the ith target (decoy) peptide is the maximal score of all PSMs matched to that peptide.

To reduce the random effects due to the random nature of our decoys we repeated the database construction 20 times with different seeds to create varied shuffled decoy peptides. We then applied TDC as well as Group-walk to the list of target-decoy scores where the latter was also given the relevant group structure as described below. We compared their power by looking at quantiles of ratios of Group-walk to TDC discovery numbers where the quantiles were taken with respect to the combined list of ratios made of 20 such ratios (one for each decoy) per each dataset. We used overall quantiles here because TDC occasionally reported 0 discoveries so taking an average of the 20 ratios per dataset would have been problematic.

#### 3.3.1 Charge and peptide length in low-resolution MS/MS data

We examined how Group-walk can take advantage of the precursor charge state and the peptide length focusing on the 7 low-resolution MS/MS data sets, because the differences in calibration were empirically pronounced there and thus served as a viable application of Group-walk. These data sets are still relevant despite the fact that they are low-resolution as the oldest one was published in 2019.

To group by charge we partitioned the peptides into groups according to the charge state of the spectrum that was maximally matched with the peptide. Note that spectra with charge states +2 or +3 include the majority of the spectra, whereas spectra with charge states +1, +4 and +5 are less frequent. The problem with having small groups is that Group-walk is more likely to exhaust them before the estimated FDR reaches the threshold *α*, thus preventing us from making any discoveries in them. To get around this, we merged the smaller groups — those that comprise less than 10% of the total number of peptides — with the previous charge state group (or, in the case of the lowest charge state group, with the subsequent group).

Grouping peptides via the length of the matched peptide sequences is more challenging given the much larger range of observed peptide lengths. However, as noted below, our simulation studies suggested that it is advantageous to partition the peptides into equally sized groups (Section 4.2). Additionally, we expect that groups of peptides with similar lengths will have a more homogenous score distribution than groups with significantly varying peptide lengths. Hence, a natural length-based partition of the peptides is to partition the range of peptide lengths into *g* roughly equal-sized groups each covering a contiguous range, or an interval of peptide lengths. Here we considered *g* = 2,3.

#### 3.3.2 Open search

Canonically, each observed spectrum is not scored against the complete target-decoy database; instead, only a subset of the peptides in the database are scored — those whose mass is within some narrow window of tolerance about the observed spectrum mass. In practice, we can extend this window as large as we like, creating what is called an *open search.* Such a search allows us to identify spectra generated by peptides that undergo modifications which significantly change their mass. Specifically, in a standard (narrow) search, the mass of the unmodified (database) peptide does not fit within the tolerance window of such a modified precursor. Such modifications of the peptide mass may occur through several mechanisms including methylation and phosphorylation [4].

After identifying the spectra using an open search we proceed as in the narrow search by assigning each target/decoy peptide a score that corresponds to the score of its maximally matched PSM. At this point we can proceed with TDC; however, in doing so we would miss out on the information that is encoded in the mass difference between the unmodified peptide and the precursor. Specifically, we expect that the mass difference for an incorrect PSM will be randomly spread over the range of possible values, whereas the mass difference of a correct PSMs will concentrate on a few values including, for example, the mass difference due to phosphorylation.

To take advantage of that information we apply Group-walk to the peptides after splitting them into two groups as follows. For each identified target/decoy peptide in the database we determine the difference between its mass and the matching spectrum’s precursor mass. We then bin the mass differences according to some regular intervals and assign the identified peptides whose corresponding mass difference falls into the top *b* most frequent intervals into one group (we used *b* = 10), and the remaining into a second group.

It is important to note that this grouping satisfies Assumption 2 required for theoretical FDR control. More specifically, knowing the mass differences between the matched peptides and the precursors (along with the winning scores and labels of correctly matched peptides), we still expect the *incorrect* matches to be independent and equally likely to be a target or a decoy win.

## 4 Results

### 4.1 Separate TDC analysis does not generally control the FDR

As motivation for creating our Group-walk procedure, we begin by demonstrating that naively applying TDC separately to each group can fail to control the FDR. In Supplementary Section 6.3 we provide the details of a simulation we carried out to illustrate this effect. Specifically, our simulations demonstrate that the actual FDR in this setup (estimated from 100K randomly drawn sets) rapidly increases beyond the FDR threshold as we increase the number of groups (Figure 1A). For example, with just three groups the actual FDR of 0.143 is already >40% higher than the nominal *α* = 0.1, and with eight groups it is >100% higher (at 0.202).

**Figure 1.**
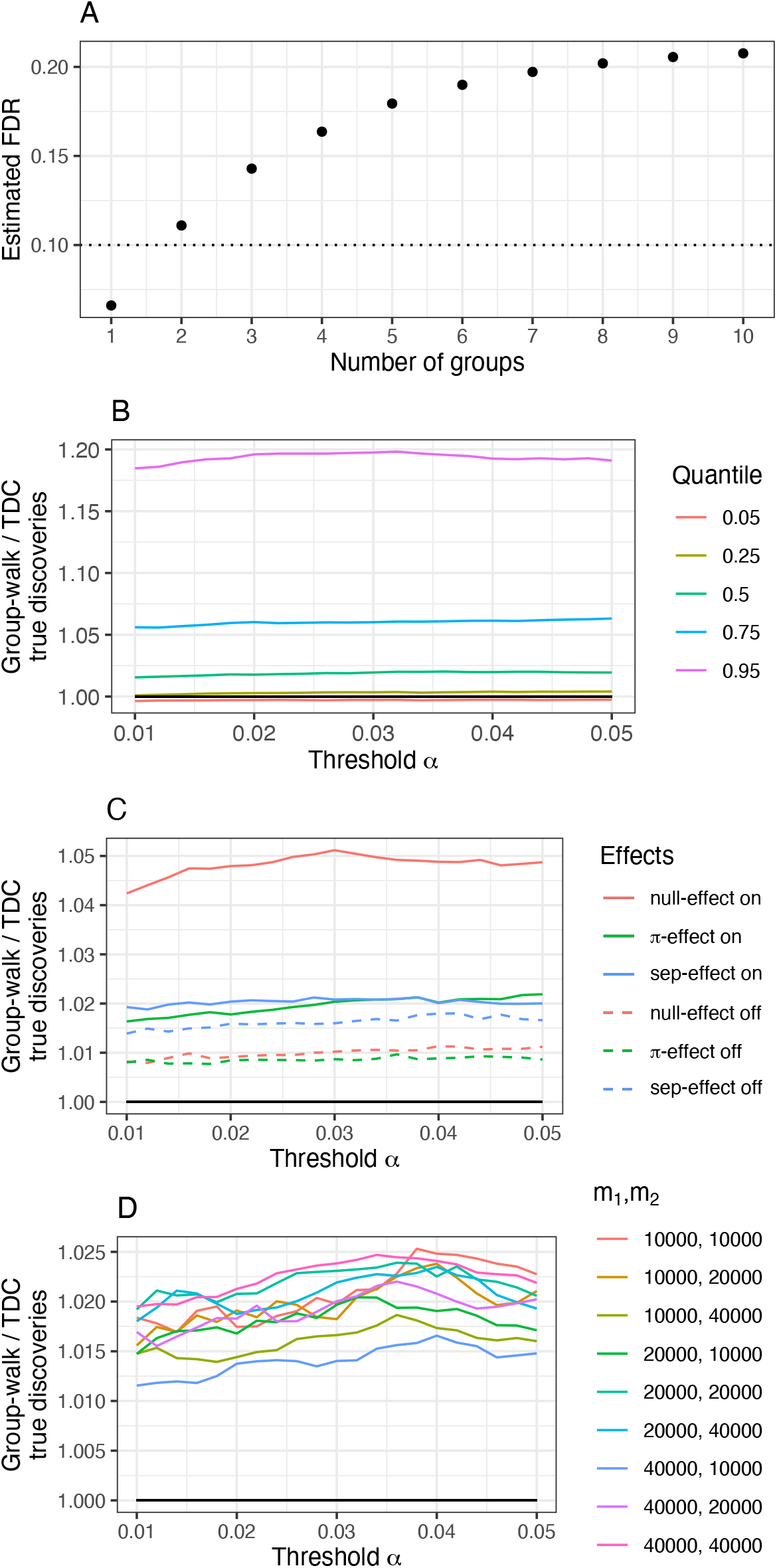
(A) The estimated FDR as a function of the number of groups. TDC is applied separately to each group and the overall FDR is averaged over 100,000 runs where the setup for each run is described in Supplementary Section 6.3. The estimated standard errors for each estimated FDR are too small to be seen in the figure. (B-D) Quantiles of the average ratio between the true discoveries from Group-walk and TDC in simulated spectrum-ID data. The average is taken over 40 runs for each of the 628 parameter combinations. (B) some primary quantiles of the 628 averaged ratios. (C) the median of the averaged ratios when we split the parameter combinations into two equal sized groups according to whether each of the three considered group effects is present (solid) or absent (dashed). (D) the median of the averaged ratio for all parameter combinations with the prescribed groups sizes (*m*_1_, *m*_2_).

### 4.2 Simulation: the spectrum-ID problem

In Figure 1B we look at quantiles of the average Group-walk-to-TDC discovery ratio using our simulated spectrum-ID scenario. The ratio’s average is taken with respect to the 40 runs, and the quantiles are with respect to the 628 parameter combinations. The plot suggests that in the context of our simulations at 1% FDR we observe a typical gain of about 1.6% more discoveries using Group-walk (median, green). That gain often grows to over 5.6% (upper quartile, blue) and occasionally to more than 18% (0.95 quantile, violet). Notably, there is typically very little loss in using Group-walk: the lower quartile still shows a very marginal gain (0.1%), and the 0.05 quantile corresponds to less than 0.4% fewer Group-walk discoveries.

In panel C we examine how Group-walk takes advantage of the three types of group effects by looking at the median of the (average) Group-walk-to-TDC discovery ratio when the data is split three times into two sets (each made of 324 parameter combinations) according to whether or not the corresponding group effect was present in the data. While Group-walk benefits from all three group-effects, the null-group effect seems to be the most prominent one in our setup.

Finally, in panel D we look at the median of the (average) Group-walk-to-TDC discovery ratio when the data is split according to the size of the two present groups (*m_i_*). Notably, Group-walk performs relatively better when *m*_1_ = *m*_2_, and its advantage decreases as the difference between the group sizes increases (blue and olive curves). This makes sense intuitively if you think about a limiting case where, say *m*_1_ ≪ *m*_2_, in which case there will be little difference between applying TDC to the combined list and to group 2 by itself.

### 4.3 Charge and peptide length in low-resolution MS/MS data

It is well understood that charge contributes to the lack of calibration for some score functions, like SEQUEST’s XCorr score [20]. Additionally, we observe a similar behaviour relating to peptide length. Here we demonstrate how the usual TDC procedure can be improved upon when applying Group-walk to such uncalibrated scores. The peptides were partitioned into 2 or more groups as described in Section 3.3.1, and again we compared the average ratio of Group-walk to TDC discoveries.

Figure 2A shows that for both charge-(2 or more groups) and length-based (2 groups) Group-walk typically offers a modest gain in power across all FDR thresholds in the considered range ([0.01,0.05]). Specifically, at the 1% FDR level the median power, when grouping by charge, increases by charge 4%, and 3.6% when grouping by length. Moreover, considering the upper quartiles we see that 25% of the time Group-walk reports at least 19% more discoveries when grouping by charge and at least 11% more when grouping by length. Examining the lower quartiles we find Group-walk still reports more discoveries than TDC: 1.4% more when grouping by charge and 0.5% more when grouping by length.

**Figure 2:**
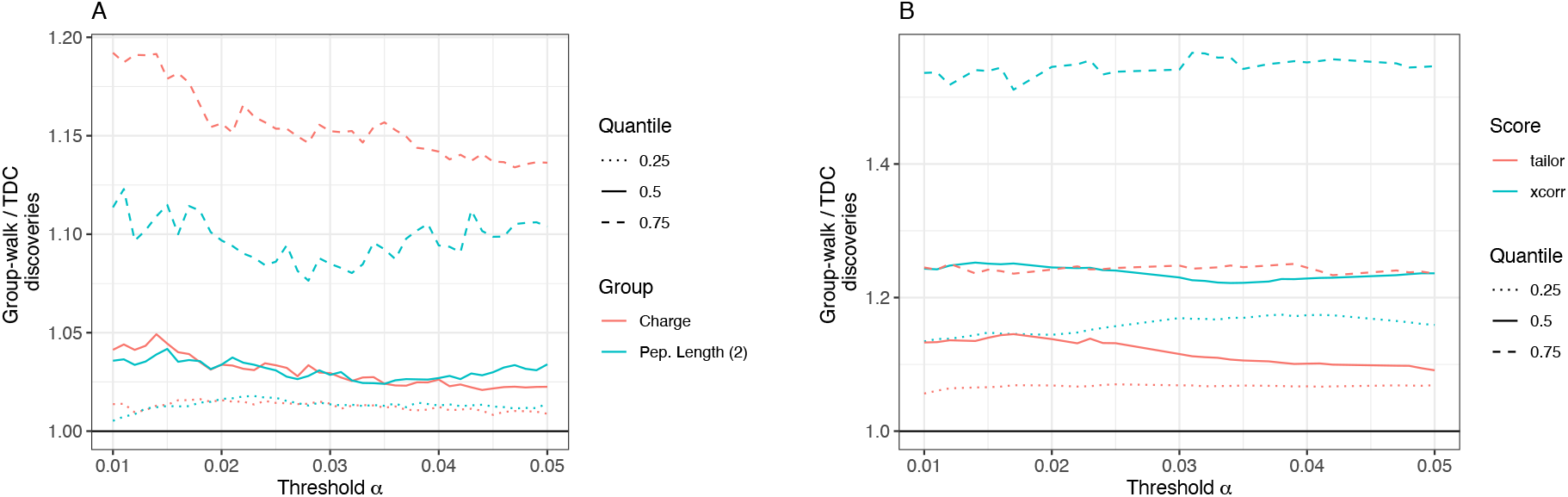
The quartiles of the ratios of Group-walk to TDC discoveries in real data. (A) Group-walk was applied to partitions of the peptides according to the charge state of the spectrum using 2 or more groups (red), as well as into 2 groups according to the length of the matching peptide (blue). (B) Results from open-searches using the XCorr score (blue) and Tailor score (red). The peptides are partitioned into two groups: the peptides whose mass difference to the spectrum’s precursor mass falls into the top *b* =10 most abundant bins in one group, and the rest in the second group.

It is worth noting that when our peptide length-based partition uses 3 groups the results are quite similar to when we partition into 2 groups as reported above (Supplementary Figure 3).

### 4.4 Open search

Comparing the performance of Group-walk and TDC in open search we see that regardless of whether the score is calibrated (Tailor) or not (XCorr) Group-walk typically delivers a fair bit more discoveries than TDC across the entire range of considered FDR thresholds (Figure 2B).

In particular, at the 1% FDR threshold Group-walk typically reports 12% more discoveries than TDC when using Tailor (solid red) and 24% more when using XCorr (solid blue). Moreover, considering the upper quartiles we see that 25% of the time Group-walk reports at least 20% more discoveries using Tailor and at least 50% more using XCorr. Examining the lower quartiles we find Group-walk still reports more discoveries: 6% more with Tailor and 15% more using XCorr.

Considering that Tailor scores are relatively well calibrated, this suggests that, as predicted, Group-walk is leveraging the fact that different groups exhibit a different proportion of correct peptide identifications and not just a difference in calibration between groups.

## 5 Discussion

TDC is the common approach to FDR control in the analysis of tandem mass spectrometry data. The implicit assumption that TDC relies on is that the data is essentially homogeneous. However, in practice the data is often made of groups of hypotheses (PSMs or peptides) with distinct group-wise characteristics. Applying TDC while ignoring this structure creates imbalanced lists of discoveries when viewed with the correct group structure in mind: the discoveries from some groups will be overly conservative while for others they will be overly liberal.

While applying a separate TDC analysis to each group addresses this problem, it might fail to control the FDR in the combined list of discoveries, and the examples we provide show that this failure can be substantial. Instead, we offer Group-walk, a novel procedure that shares some of the characteristics of separate analyses while still controlling the combined FDR. It is worth noting that, particularly when using the common 1% FDR threshold, Group-walk can possibly deliver even more discoveries than the, non-rigorous, separate analyses. Indeed, the +1 “penalty” in the estimated FDR is applied to each group in the latter case, but Group-walk uses a combined FDR estimate so in its case there is a single penalty.

Group-walk can be considered as a specialization of a slightly modified version of AdaPT, which is a recently published general framework for the problem of multiple hypothesis testing with generic side information [22]. Note that as a general framework AdaPT does not specify how to advance its analogue of what we call the front, leaving it instead as a generic function called `UPDATE` (a specialized version of `UPDATE` is offered in [22], but it only applies for the classical p-value setup rather than the competition-based setup we study here). Still, showing that Group-walk is consistent with the general requirements of AdaPT’s `UPDATE` allows us to establish that Group-walk controls the FDR in the finite sample setting.

To be effective, Group-walk requires typically at least K hypotheses in each group (throughout this work we used *K* = 40). Some other approaches in the spirit of separate TDC analyses have been offered for cases where some groups are small, but no finite-sample FDR control was established (e.g. [13]).

We demonstrated using simulated data how Group-walk can benefit from the varying types of group structures: null-effect, separation-effect, and *π*-effect. It should be noted that both the separation- and *π*-effects have general trends that affect how well Group-walk can take advantage of them. For example, when *μ* is large enough so the alternative distribution becomes sufficiently separated from the null distribution, the problem becomes trivial, and TDC will do just as well as Group-walk even if the data exhibits a significant group *π*-effect.

This raises the important question of how do we know when should we use Group-walk. For now we leave this for future research, noting that our results show that the penalty of using Group-walk in the absence of group structure seems rather marginal, particularly when compared with the significant gains it potentially offers.

A particularly appealing property of Group-walk is that it frees us from worrying about whether our score is sufficiently calibrated with respect to some property of the data, such as the precursor’s charge state, or put differently, whether our data exhibits a group null-effect. Indeed, as we saw in the discovery gains Group-walk offered when grouping the low-resolution PRIDE datasets by charge, all we have to do is partition the data into groups according to the considered feature and apply Group-walk which will then take advantage of the null-effect.

While we also saw gains when analyzing the PRIDE data sets according to the peptide length and to the mass difference in open search, we note that it is not always obvious how to partition the hypotheses into groups. This is even more the case if the feature we consider is not discrete. This suggests another promising direction for future research, which is to go back to AdaPT’s view of having more general side information rather than just a group ID. Note that again, it is not obvious what needs to be done, as such an approach would still require specializing AdaPT’s generic UPDATE function for our competition-based setup.

While TDC has been in use for some time now in computational mass spectrometry, Barber and Candés recently used the same principle in their knockoff+ procedure to control the FDR in feature selection in a classical linear regression model [2]. Moreover, following their work and the introduction of a more flexible formulation of the variable selection problem in the model-X framework of Candés et al. [5], competition-based FDR control has gained a lot of interest in the statistical and machine learning communities. Knockoffs and decoys serve the same purpose in competition-based FDR control. In particular, Group-walk is applicable to the knockoff context just as well as it is to TDC. That said, in the kind of feature selection problems considered by Barber and Candés the fraction of false nulls (relevant features) is often too low to effectively employ Group-walk.

## 6 Supplementary Material

**Algorithm 1:**
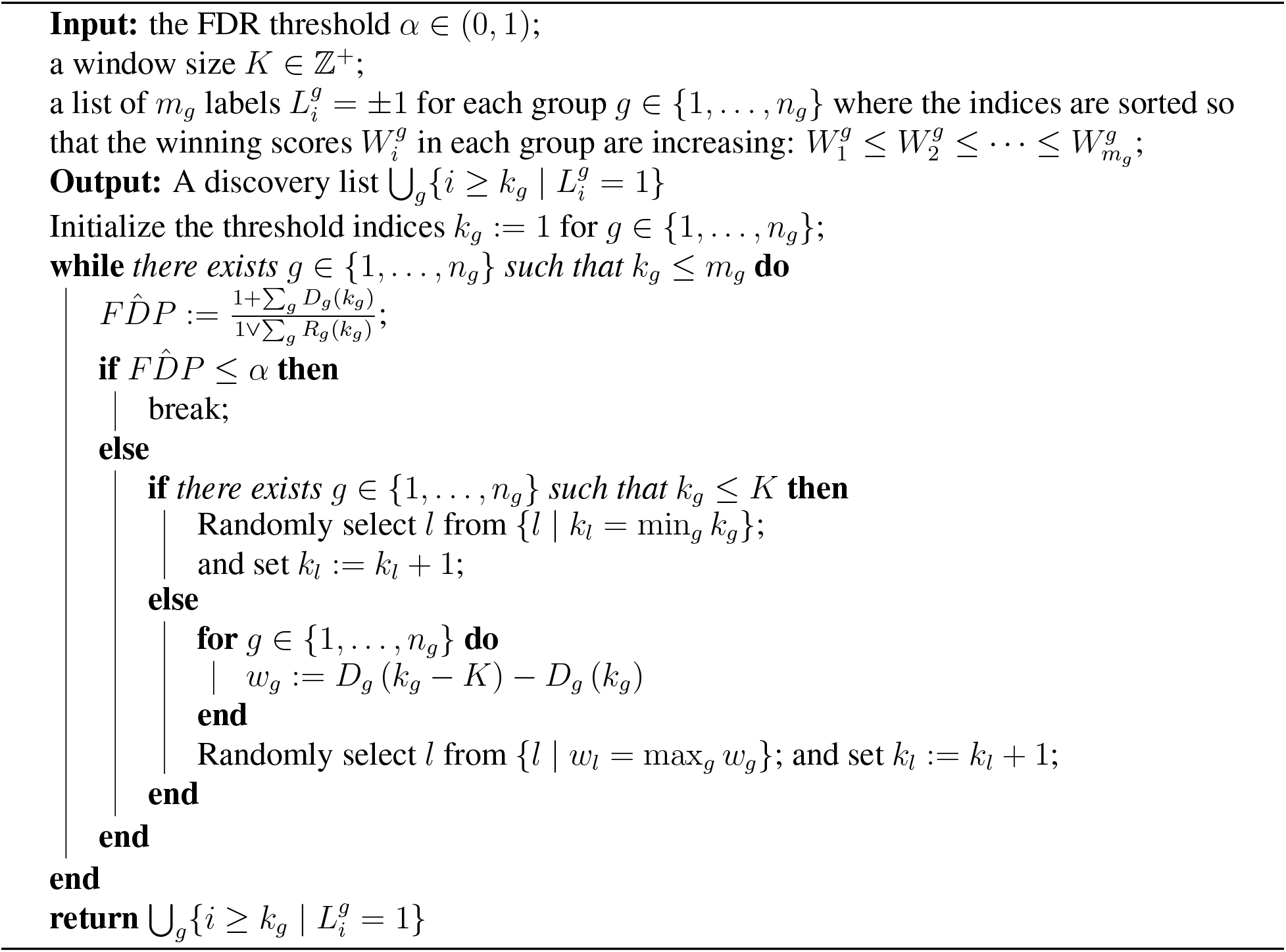
Group-walk.

### 6.1 Simulation: the spectrum-ID model in detail

We associate with each (virtual) “spectrum”, *σ_i_*, a score *X_i_* as follows. If *σ_i_* is native (cf. Section 2) then *X_i_* models the match between *σ_i_* and the peptide that generated it, so we draw *X_i_* ~ 1 – Beta (*a, b*), where we fix *a* = 0.05 and we consider b as the *separation parameter* between true and false null scores. Otherwise, *σ_i_* is foreign so we set *X_i_* to the lowest possible score which is *X_i_* = 0 here. All the draws described here are done independently.

In addition, assuming that associated with each spectrum are *n* candidate peptides (those within the tolerance of the measured precursor mass, here we used *n* = 100), we associate with each native spectrum a score *Y_i_* corresponding to the best match to *σ_i_* among the remaining *n* – 1 candidate target peptides, whereas for a foreign spectrum *Y_i_* corresponds to the best match among all n candidates. As all these target matches are considered random we model *Y_i_* as the maximum of *n* – 1 (or *n*) Unif (0,1) scores, i.e., we sample *Y_i_* ~ Beta (*n* – 1,1) (*Y_i_* ~ Beta (*n*, 1)). The target PSM score is then *Z_i_* = *X_i_* ∨ *Y_i_*.

We also associate with each spectrum a decoy PSM score, 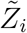, modeling the best match to *σ_i_* among *n* candidate decoy peptides. Consistent with our Unif (0,1) model of a random match we therefore draw 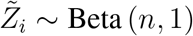. It follows that Assumptions 1 and 2 are only approximately valid under this particular model: for native spectra there is a slightly larger chance a true null will be a decoy win. As this creates a slightly conservative bias it is not concerning as far as FDR control. Additionally, if *σ_i_* is foreign then *Z_i_* ~ Beta (*n*, 1) so for a foreign spectrum target and decoy wins are equally likely.

To make the score uncalibrated, we define the winning scores as 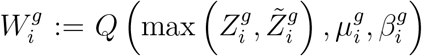, where *Q* (·, *μ, β*) is the quantile function of the Gumbel distribution with parameters *μ, β*. The latter location and scale parameters are sampled from a population of such parameters generated by fitting Gumbel distributions to PSM scores generated from real yeast data, with a presumed charge state of 2 or 3. Further details can be found in [21].

Our simulated dataset of PSM scores consists of two groups (*n_g_* = 2) where each group has *m_g_* null hypotheses (“the PSM is incorrect, i.e., 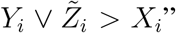). A fraction of *π_g_* of the target scores 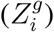 within each group, corresponding to the native spectra, are sampled as described above from the alternative distribution (*X_i_* ~ 1 – Beta (*a, b*)), and the remaining target scores, as well as all the decoy scores 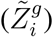, are sampled from the null distribution. All the draws are made independently.

We varied the group sizes *m*_1_, *m*_2_ ∈ {1e4, 2e4, 4e4}, the separation parameters *b*_1_, *b*_2_ ∈ {5, 50} and the proportion of native spectra in each group *π*_1_, *π*_2_ ∈ {0.3, 0.6, 0.9}. In addition, to study the group null-effect we create two different types of datasets. In the first type the Gumbel parameters that induce the non-calibrated feature of the data, 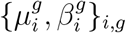 are all randomly drawn with replacement from the same population of location and scale parameters. Any such homogeneous dataset is uncalibrated but there is no group null-effect. The second data type is generated where the Gumbel parameters of the first group are drawn from the population of parameters that were originally fitted to PSMs with a presumed charge state of 3, while the Gumbel parameters of the second group are drawn from the set fitted to a presumed charge of 2. As noted in the Introduction, different charge states are known to be associated with a group null-effect.

Again, we generated 40 samples for each combination of parameters and examined the average ratio of Group-walk (K = 40) to TDC true discoveries number for each of the 628 distinct parameter combinations.

### 6.2 Supplementary Analysis of mass spectra

The following data sets that we used were sampled from projects available from the Proteomics Identifications Database (PRIDE) [28]. One spectrum file was randomly selected and downloaded from each of 14 randomly selected projects such that 7 were high-resolution spectra files and 7 were low-resolution spectra files. Each project was submitted to PRIDE no earlier than 2018. The 7 low-resolution spectrum files came from PXD012522, PXD013482, PXD014774, PXD015485, PXD017622, PXD022092 and PXD027867. The 7 high-resolution spectrum files came from PXD006856, PXD008996, PXD012611, PXD013274, PXD019186, PXD022257 and PXD025130. We also downloaded the target peptide database from PRIDE except for human data where we used the UniProt database (UP000005640) downloaded on 9/11/2021.

We used the Crux toolkit [27] to assign peptides to spectra as follows. First, for each PRIDE dataset, the Crux command tide-index [7] was used to construct the concatenated target-decoy peptide database with the (Param-Medic) option `--auto-modifications-spectra` to account for variable modifications [23]. To reduce the random effects due to the random nature of our decoys we repeated the database construction 20 times with different seeds to create varied shuffled decoy peptides, which our results were aggregated over. Tide-search [7], with parameter settings optimized by Param-Medic [24], was then used to generate the PSM scores by searching the concatenated target-decoy peptide database with the exact options as follows:

- When grouping by the precursor charge state and the peptide length we used the following options in tide-search: `--auto-precursor-window warn, --auto-mz-bin -width warn, --concat T`.
- In the open search analysis we used: `--auto-mz-bin-width warn, --precursorwindow-type mass,--precursor-window 100,--use-tailor-calibration T, --concat T`. In addition, we used both the XCorr score, and a calibrated version of XCorr score called *Tailor* score [30]. Note, that the option `--use-tailor-calibration T` returns both scores.

We restricted the analysis of grouping by charge and peptide length to the 7 low-resolution datasets whereas the open search analysis was done on all 14 datasets. To discretize the mass difference in the latter analysis we used multiples of the canonical value λ = 1.0005079 Da, as the breaks that define the intervals/bins of mass differences: {…, —3λ/2, —λ/2, λ/2, 3λ/2,… }.

### 6.3 Separate TDC analysis does not generally control the FDR

It is not difficult to find examples showing separate TDC analyses fail to control the overall FDR in a finite sample setting. For example, consider the following setup for two groups. For *g* ∈ {1.2}, let *m_g_* be the number of hypotheses in group *g* and *π_g_* be the fraction of false nulls. We set *m*_1_ = 100, *m*_2_ = 1000 and *π*_1_ = *π*_2_ = 0.1. With 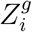 and 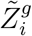 denoting the target and decoy scores of the ith hypothesis in group *g*, respectively, we draw all scores independently as follows: 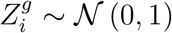 if 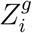 comes from a null hypothesis, and 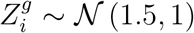 if 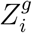 comes from a false null hypothesis. The decoy score 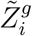 is distributed 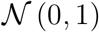. For each hypothesis, we determine the winning scores and labels as defined in Section 2 and apply the TDC procedure separately, resulting in two discovery lists. Repeating the above simulation for 100,000 runs at an FDR threshold of *α* = 0.25, we found the *combined* empirical FDR was 0.258 with an estimated standard error of 0.0005; 16 standard errors above the threshold.

While the above failure to control the FDR is statistically significant, it is rather negligible in terms of the magnitude of the excess FDR. However, by increasing the number of groups we can readily construct examples where the actual FDR of separate TDC analyses is substantially higher than the nominal threshold.

Specifically, we consider *n_g_* independent and identically distributed groups where each group is made of 22 hypotheses, 6 of which are true null hypotheses and 16 are false nulls. Moreover, we assume the range of possible scores for each hypothesis is such that the order of the hypotheses is deterministic. Specifically, when sorted in increasing order, first come 3 of the null hypotheses followed by 8 of the false nulls, then the remaining 3 nulls followed by the remaining 8 false nulls. For each false null hypothesis we assume the target always wins, whereas the labels of all true null hypotheses are independently and uniformly drawn.

We repeatedly and independently drew the sequence of labels for each of the *n_g_* groups, applied TDC separately to each group at level *α* = 0.1 and noted the actual FDP in the combined list of discoveries. We estimated the FDR in the combined list by averaging the actual FDP over 100,000 sets of draws for each value of *n_g_*, as we increased it from 1 to 10. These estimated FDR values given in Figure 1A demonstrate how rapidly the excess FDR grows in this case, indicating that even for a small number of groups, separate analyses can provide a poor method of FDR control. For example, with just 3 groups at 0.143 the actual FDR is already over 40% higher than the nominal *α* = 0.1 and with 8 groups at 0.202 it is more than 100% higher.

Let *V_g_* denote the number of true null target wins discovered by TDC in group *g*, and let *R_g_* be the corresponding number of all discoveries. It is not hard to show that as *n_g_* → ∞ the FDR of the separate TDC analyses will converge to the mFDR of each group which is defined as *E*(*V_g_*)/*E*(*R_g_*). Given the size of the problem, it is straightforward to evaluate the ratio of expectations and find that in this case mFDR = 1.15625/5.40625 ≈ 0.214; significantly over the nominal 0.1.

### 6.4 Group-walk is a specialized version of AdaPT

We establish Group-walk’s FDR control by showing it is a special case of (a slightly modified version of) the Adaptive p-value Thresholding (AdaPT) framework of Lei and Fithian. AdaPT was developed to address the problem where associated with each hypothesis is a p-value *p_i_*, as well as a predictor *x_i_* encoding contextual information about the hypothesis [22]. In our case the p-value is the 1-bit p-value (1/2 for a target win and 1 for a decoy win), and *x_i_* = (*W_i_, G_i_*).

AdaPT progresses through iteratively defined rejection thresholds, *s_t_*(*x*), that are analogous to Group-walk’s front. Indeed, *s_t_* describes the front position at step *t* of the procedure and for each fixed *x* the rejection thresholds are monotone decreasing in *t*. To show that Group-walk is an adaptation of AdaPT we introduce t as the same kind of step counter to Group-walk, and we define *s_t_*(*x*) in that case as 0 for all the hypotheses that Group-walk’s front has passed over by step *t*, and as 1/2 for all the other hypotheses: with **k**_*t*_ = (*k*_1_,…, *k_n_g__*) denoting Group-walk’s front at step *t*, 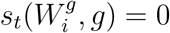 for all 1 ≤ *i* < *k_g_* and 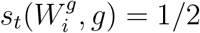 for all *k_g_* ≤ *i* ≤ *m_g_*.

With this definition of *s_t_*, and provided we slightly modify the definition of AdaPT’s *A_t_* to use a strict inequality:

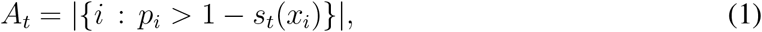

A_t_ coincides with Group-walk’s 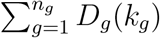. Similarly AdaPT’s 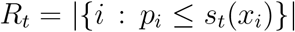 coincides with Group-walk’s 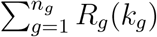. It follows that with this modification their estimated FDR, (1 + *A_t_*)/(1 ∨ *R_t_*), coincides with Group-walk’s and hence AdaPT’s rejection criterion coincides with Group-walk’s in this case.

AdaPT does not specify how to adjust its front *s_t_* except to note that the update needs to be a function of *A_t_, R_t_, s_t_* as well as of all the contextual information ((*W_i_, G_i_*) in our case), and all its 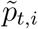 which are defined in (2) of [22]. Here is where we need to make our second change to AdaPT’s original definitions. Specifically we change the second inequality in their definition (2) to a weak one:

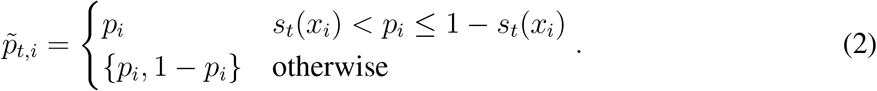

With the latter change it is easy to see 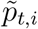 resolves the target/decoy label, *L_i_*, for any hypothesis that the front has passed over by step *t* and the label remains unresolved otherwise. It follows that Group-walk’s update procedure, which moves the front by moving one of the indices forward, *k_g_*: = *k_g_* + 1, based on the information gathered from the labels of hypotheses the front already passed over, as well as on the scores and the group assignments is consistent with AdaPT’s generic update protocol.

Finally, we verified that Theorem 1 of [22], which proves that AdaPT controls the FDR in the finite sample setting, carries through with our two modifications above: (1) and (2). It follows that under Assumption 2 Group-walk also controls the FDR in the finite sample setting.

### 6.5 Supplementary Figures

**Figure 3:**
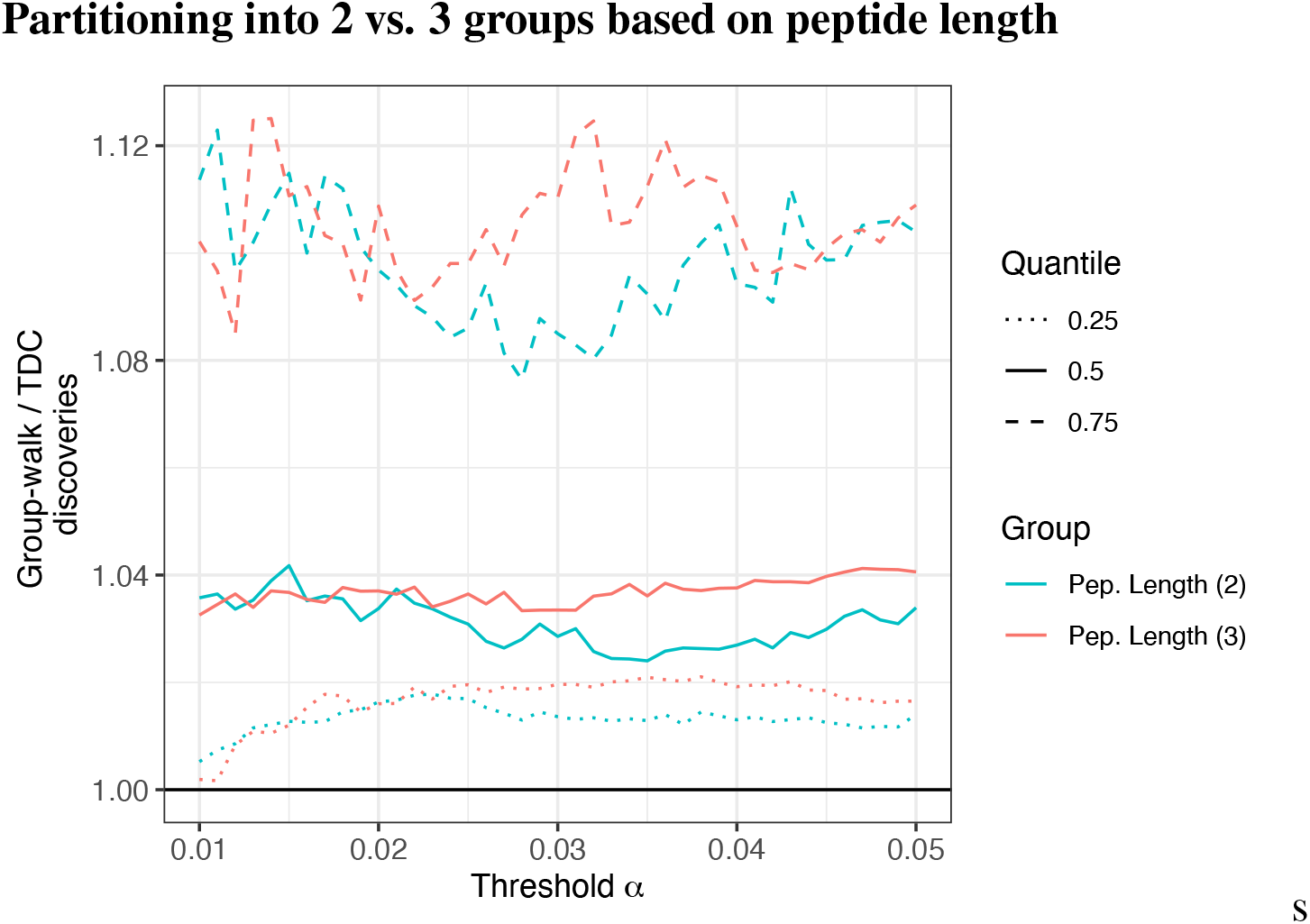
The figure presents the quartiles of the ratios of Group-walk to TDC discoveries when partitioning the peptides via the the length of the matching peptide into either 2 (blue) or 3 (red) groups.

